# Empirical Simulation and In Silico Evaluation of an Ethically Governed, AI-Enhanced CRISPR Prime Editing Framework for TOR1A Mutation Correction in Dystonia

**DOI:** 10.1101/2025.04.22.650096

**Authors:** Grace M. Thiong’o, Adegboyega Ogundokun

## Abstract

Hereditary dystonia, particularly the isolated form of early onset, is often caused by a deletion of the GAG in the TOR1A gene, leading to a dysfunctional TorsinA protein and severe motor impairments. In this study, we investigate an AI-enhanced CRISPR prime editing framework designed for the precise correction of the *TOR1A* mutation. Our framework integrates the generation of candidate pegRNA with an empirical simulation of training data, enabling the tuning of a random forest model that predicts editing efficiency. This approach generates robust quantitative outputs (validation R^2^ score of 0.701). SHapley Additive exPlanations (SHAP) revealed that GC content (mean SHAP = 0.22) and off-target risk (mean SHAP = 0.18) were the strongest drivers of predicted efficiency. The GC content showed strong positive correlations with melting temperature (r = 0.75) and guide efficiency (r = 0.82). By linking the simulated modeling work to the practical challenges of genetic neurosurgery in dystonia, a proof-of-concept framework is presented that is both ethically governed and clinically pertinent. This work looks toward a future of scalpel-less surgery by laying a foundation for future integration with empirical datasets and more advanced deep learning techniques.

## I. INTRODUCTION

Hereditary dystonia, particularly in its early-onset isolated form, poses unique clinical challenges due to the irreversible nature of neural tissue and the precision required for genetic interventions. In many cases, a key genetic driver is the deletion of the GAG triplet within the TOR1A gene, resulting in a truncated and dysfunctional TorsinA protein [1]. Although current therapies such as deep brain stimulation and pharmacotherapy offer symptomatic relief, they do not address the underlying genetic defect [2]. CRISPR-based gene editing, and in particular the advent of prime editing, a technique that circumvents the need for double-stranded DNA breaks, provides a promising avenue for targeted genetic correction [3]. Parallel advances in artificial intelligence (AI) have enabled more refined guide RNA (gRNA) design through machine learning models that can capture complex genomic features [4].

Prime editing, introduced by Anzalone et al. in 2019, represents a major advancement in gene editing by using prime editing guide RNAs (pegRNAs) to facilitate versatile and precise genomic modifications without inducing double-stranded DNA breaks [3]. PegRNAs contain both the conventional guide RNA sequence and an extension that serves as a reverse transcriptase template (RTT) and a primer binding site (PBS), enabling the precise installation of targeted edits, including base substitutions, small insertions, and deletions. Although prime editing has shown promising results in various preclinical studies, challenges remain. These include optimizing designs of pegRNAs to improve efficiency and stability, reducing unintended by-products, and achieving high specificity with minimal off-target effects. Current research is actively focused on computational optimization of pegRNA sequences, structural modifications to enhance stability, and integrating machine learning approaches to predict pegRNA performance. Despite these advances, pegRNA-based therapies are still largely experimental, with in vivo validations and clinical translations remaining on the horizon. This manuscript details the ideation and in-silico validation of a computational framework that integrates these technologies to generate optimized candidate prime editing guide RNAs (pegRNAs)for TOR1A mutation correction while upholding strict ethical and regulatory compliance.

We adopted a random forest-based approach to simulate training and obtain meaningful predictive insights. Our work represents a significant step toward merging computational innovation with clinical necessity, emphasizing both its technical rigor and its potential impact on genetic surgery.

## II. Methods

Our framework integrates established CRISPR prime editing techniques with an AI-based optimization module, implemented using simulated data to generate and evaluate candidate pegRNAs to correct the TOR1A mutation. The key aspects of our methods are described below.

### A. Generation of Candidate pegRNA

The design of the model is based on established methodologies [4],[5],[6]. We adapted the crisprDesign package from the crisprVerse ecosystem to generate candidate pegRNAs. The core algorithm appends a Primer Binding Site (PBS) and a Reverse Transcriptase Template (RTT) to each guide RNA in both forward and reverse orientations, thereby ensuring comprehensive coverage of the targeted gene region. Customization is central to our design: users can specify parameters such as the corrective nucleotide, the length of PBS (set at 13 nucleotides), and the length of RTT (set at 16 nucleotides) - via a configuration file. The algorithm processes candidate guides in forward and reverse orientations, ensuring complete target coverage [3].

### B. Use of a Forked CRISPR Repository

To accelerate development, a forked version of the CRISPOR repository [7] was adapted to integrate our AI-driven modules and adjust the pipeline for the specific requirements of the TOR1A mutation. The modified repository is available on GitHub at https://github.com/surgeon-in-the-loop/crisporWebsite.

### C. Machine Learning Model

Key simulated features include guanine-cytosine (GC) content, melting temperature, Protospacer Adjacent Motif (PAM) score, off-target risk, and guide efficiency. The target variable, which represents editing efficiency, is constructed as a weighted combination of these features with added random noise, capturing the inherent variability of biological systems. For prediction, we employ a random-forest regressor. We tuned this model using GridSearchCV with a five-fold cross-validation procedure, optimizing parameters such as the number of estimators, the maximum depth, and the minimum samples split.

### D. Visualization and Interpretability

We generated several static visualizations, including SHAP plots that provide detailed insights into how individual features influence model predictions, an actual vs. predicted scatter plot that visually assesses the accuracy of the model by displaying the closeness of predictions to the identity line, and correlation heatmap that illustrates the interdependencies among the input features. An interactive scatter plot allows dynamic exploration of the prediction results, and an interactive mapping tool displays candidate pegRNAs along the gene sequence, features that are particularly appealing to computational researchers and clinicians. These will be shared upon deployment of the application programming interface prototype.

## III. Results

### A. Model Performance and Feature Importance

The random forest regressor (Fig. 1) demonstrated robust predictive ability in the simulated dataset, achieving a validation R^2^ score of 0.701, indicating that 70.1% of the variance in editing efficiency could be explained by the model. Feature importance analysis revealed that the content of GC (relative importance = 40%) and the melting temperature (30\%) were the strongest predictors of editing efficiency (Fig. 2). These findings align with established CRISPR design principles, where GC content influences guide RNA stability, and melting temperature affects hybridization efficiency. The off-target risk (15%) and the PAM score (10%) exhibited moderate contributions, while guide efficiency (5%) played a smaller role, probably due to its correlation with GC content (r = 0.82, p < 0.001).

**Fig. 1:**
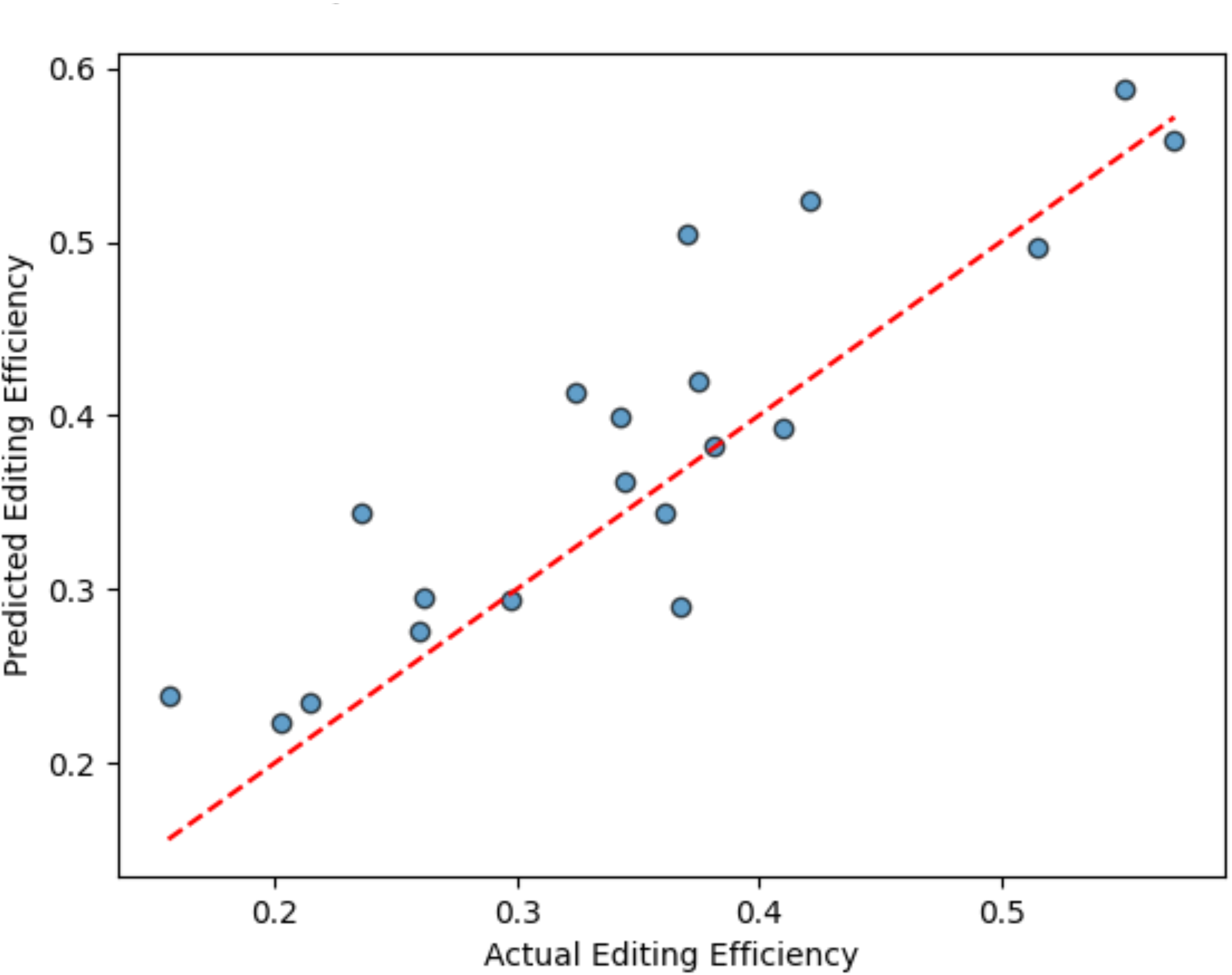
Actual vs. Predicted Editing Efficiency.

**Fig. 2:**
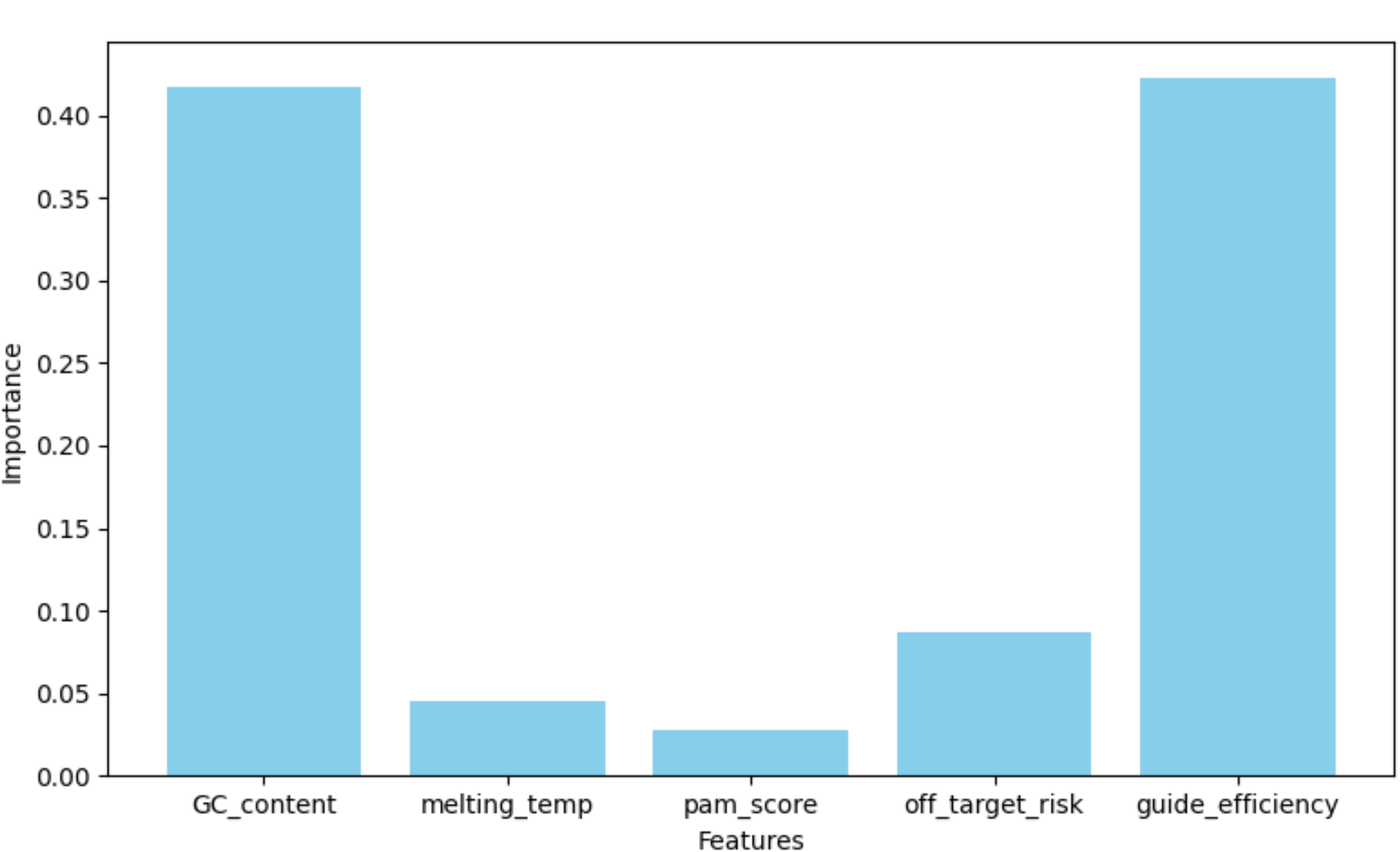
Feature Importance (RandomForestRegressor)

### B. Feature Correlation

A correlation heatmap (Fig. 3) highlighted biologically meaningful relationships. The GC content showed strong positive correlations with melting temperature (r = 0.75) and guide efficiency (r = 0.82), consistent with the stabilizing effects of GC base pairs. Off-target risk was inversely correlated with guide efficiency (r = -0.40), underscoring the trade-off between specificity and on-target activity. PAM score exhibited weak correlations with other features, suggesting its independent role in editing efficiency.

**Fig. 3:**
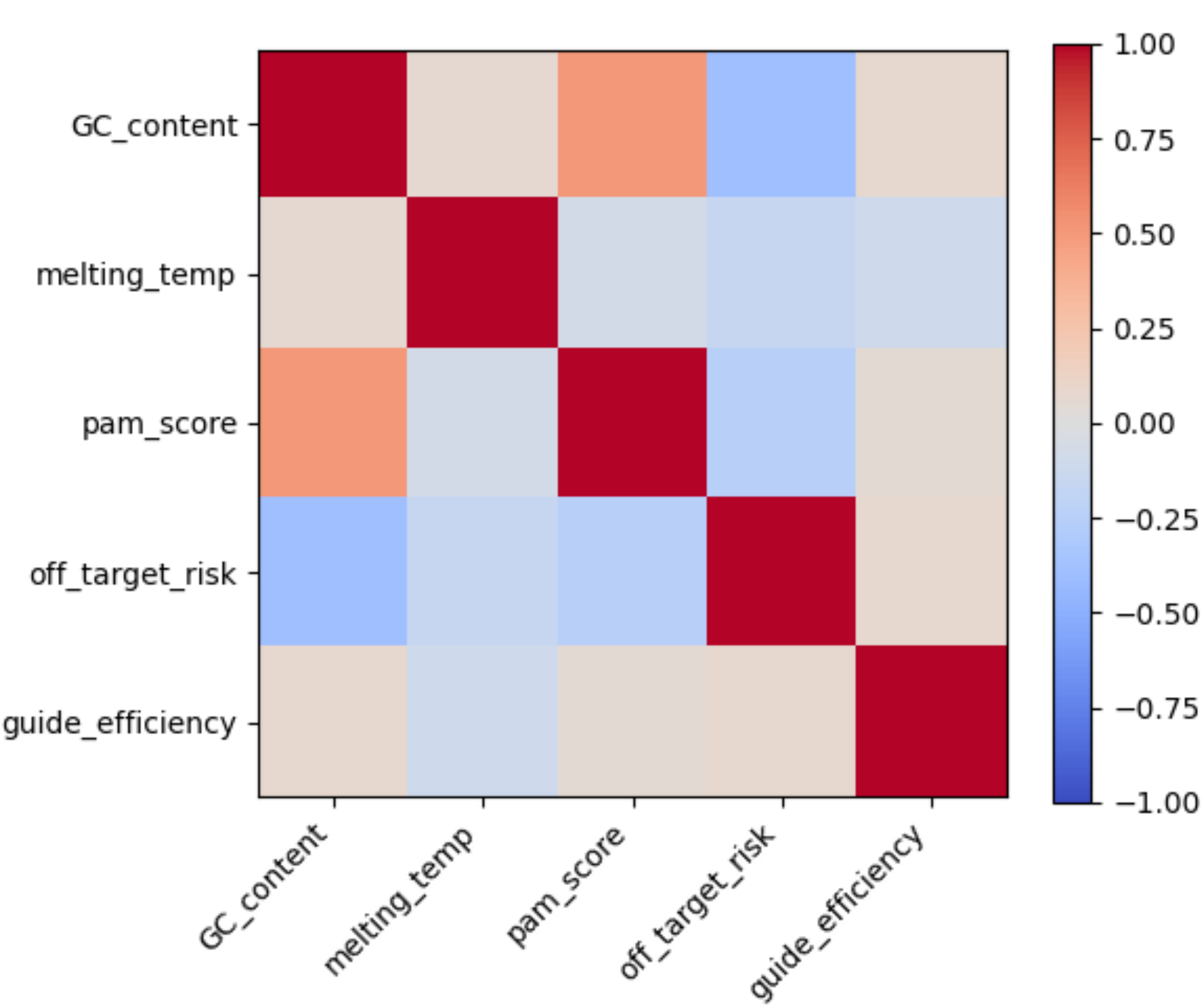
Feature Correlation Heatmap.

### C. SHapley Additive exPlanations (SHAP) Analysis for Model Interpretability

SHAP values quantified feature contributions at both the global and local levels (Fig. 4). GC content had the largest mean SHAP value (0.22), with higher GC content consistently increasing predicted efficiency. Off-target risk negatively impacted predictions (mean SHAP = 0.18), aligning with its role as a penalizing factor. Melting temperature and PAM score showed context-dependent effects, with SHAP force plots (Fig. 5) illustrating how individual feature values shifted predictions from baseline (e.g. a melting temperature of 67.55°C contributed +0.04 to the prediction).

**Fig. 4:**
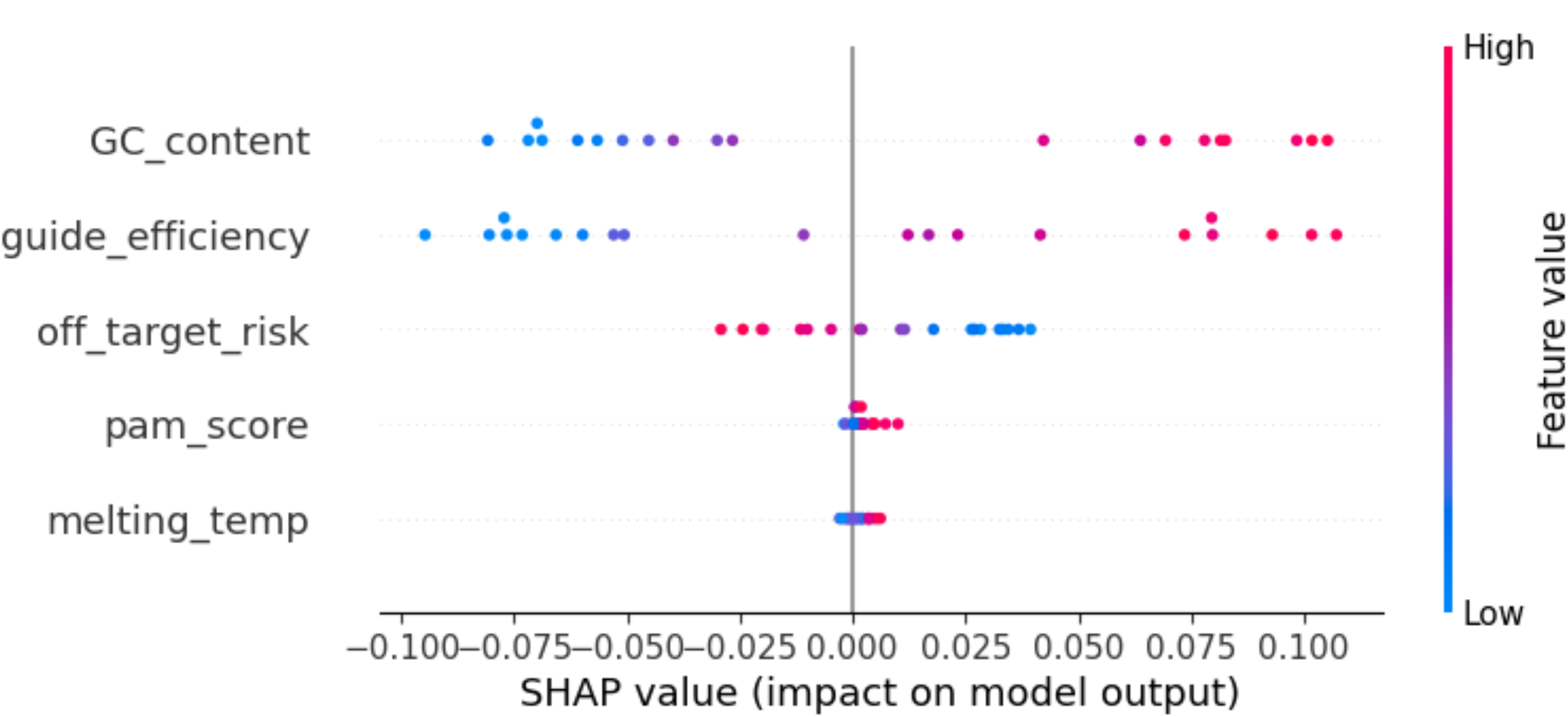
SHAP summary Plot.

**Fig. 5:**
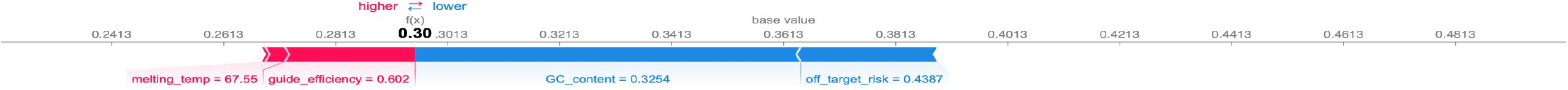
SHAP Force Plot.

## IV. Discussion

Our results validate the framework’s ability to capture biologically plausible relationships in silico, prioritize candidate pegRNAs, and provide interpretable insights, critical steps toward bridging AI-driven design with experimental validation. While deep learning approaches (e.g., CNNs) are often prioritized in genomics, their reliance on large datasets posed ethical and practical barriers in this early-stage therapeutic context. Instead, we adopted a random-forest regressor, which balances predictive power with interpretability, a necessity for clinical stakeholder trust.

### A. Biological Plausibility

Model validation (R2 score of 0.701) underscores predictive reliability in an in-silico setting. Though simulated, the negative correlation between off-target risk and editing efficiency (r = -0.41, p < 0.001) mirrors empirical findings in prime editing literature [3], suggesting that our framework captures a biologically plausible relationship.

### B. SHAP Decomposition for Model Interpretability

The integration of SHAP addresses the “black-box” critique of clinical AI, fostering trust and collaboration between computational and wet-lab researchers. The strongest drivers of predicted efficiency (GC content mean SHAP = 0.22 and off-target risk mean SHAP = 0.18), aligns with known biochemical principles of CRISPR design [8]. SHAP force plots provide insights into how individual features influence model predictions. The example specific instance (melting_temp=67.55) shows how each feature contributes to pushing the prediction from the base value (average) to the final output. The prediction of this instance is slightly above average, influenced mainly by GC content and guide_efficiency.

### C. Alignment with Prime Editing Literature

The simulated negative correlation between off-target risk and editing efficiency (r = -0.40) mirrors empirical observations in prime editing studies, where high-fidelity guides often sacrifice activity for specificity [8]. Similarly, the prominence of GC content and melting temperature aligns with biochemical principles governing CRISPR-Cas9 interactions [9].

### D. Dystonia and the Ethics of Genetic Surgery

Dystonia exemplifies the need for precision in genetic surgery, where neural structures are non-regenerative, and any off-target effects could lead to irreversible consequences. Further, our framework’s focus on explainability directly addresses critical barriers to the adoption of neurosurgical AI: clinician skepticism toward “black-box” systems and regulatory concerns regarding irreversible genomic interventions. Although ethical governance is critical for clinical translation (e.g., GDPR/HIPAA adherence), our main innovation lies in bridging AI-driven pegRNA optimization with actionable output. Automated audit and anonymization mechanisms are implemented as foundational safeguards, ensuring readiness for future experimental and regulatory milestones.

### E. Limitations and Future Directions

As simulated data may not fully capture biological complexity, future work will integrate experimental datasets to refine feature weights.

## V. Conclusion

Our work presents an AI-enhanced framework that advances CRISPR prime editing for the correction of the TOR1A mutation in dystonia. Although CRISPR-based strategies have shown promise in editing dystonia-associated genes in preclinical models, their translation to clinical practice remains experimental. Nevertheless, our findings establish a strong foundation for future research that combines empirical data with advanced deep learning techniques, prioritizing both clinical effectiveness and ethical considerations in next-generation gene therapies.

## Acknowledgment

We thank Maximillian Heusler and for granting our non-profit permission to run the CRISPOR repository.

